# Architecture of an asymmetric mycobacterial short chain/long chain acyl-CoA carboxylase

**DOI:** 10.1101/2025.11.15.688623

**Authors:** Edukondalu Mullapudi, Hai Minh Thai, Luiz Pedro Sório de Carvalho, Matthias Wilmanns

**Affiliations:** European Molecular Biology Laboratory, Hamburg Unit, 22607 Hamburg, Germany; Department of Chemistry, The Herbert Wertheim UF Scripps Institute for Biomedical Innovation & Technology, FL 33458, USA; The Skaggs Graduate School, The Scripps Research Institute, La Jolla, CA 92037, USA; University Medical Center Hamburg-Eppendorf, 20246 Hamburg, Germany

## Abstract

Endogenous extraction has revealed a mycobacterial hybrid acyl-CoA carboxylase (ACCase) complex exhibiting distinct long-chain (LC) and short-chain (LC) acyl-CoA carboxyl transferase (CT) activities. The presence of two different CT subunits (AccD4, AccD5) is triggered by a unique AccE5 dimer. AccE5 also generates a flexible biotin carboxylase (BC) / CT arrangement through a 9-stranded β−barrel, which rotates the entire BC assembly by ∼90° in the presence of acyl-CoA substrates. These data demonstrate that asymmetric, multi-substrate ACCases differ fundamentally from symmetric, single-substrate ACCases.

## Main

*Mycobacteria* possess a complex cell envelope enriched with mycolic acids that are critical for bacterial survival and pathogenicity. ACCases are multienzyme complexes that catalyze the ATP-dependent carboxylation of acyl-CoA derivatives of variable lengths to generate precursors of mycolic acids and other lipids (1–3). Because of their functional roles and structural properties, mycobacterial ACCases are well established drug targets (4). ACCases generally consist of BC and CT subunits within higher oligomeric assemblies, in which BC-linked biotin carboxyl carrier protein (BCCP) domains transfer BC-produced carboxybiotin to the CT active site for acyl-CoA carboxylation. Many mycobacteria contain an expanded repertoire of three BC subunits, up to six CT subunits, and a unique ε-subunit (AccE5) of unknown function (1, 5, 6). While mycobacterial AccD1/AccA1 and AccD2/AccA2 assemblies have been shown to form symmetric holoenzymes with single-substrate specificity (1), any mechanism on how the other ACCase protein components assemble has remained unknown.

To uncover this conundrum, we purified endogenous ACCase complexes from *Mycobacterium smegmatis.* Among the previously established AccD1/AccA1 and AccD2/AccA2 dodecameric assemblies (3), we discovered a previously unknown ACCase complex, in which we identified the presence of one distinct BC subunit (AccA3), two different CT subunits (AccD4, AccD5) and the AccE5 subunit, which is absent in other ACCase complexes **(Extended Data Fig. 1a,b).** Consistent with the presence of two different CT subunits, this hybrid complex exhibited catalytic activity for both SC (acetyl-CoA, propionyl-CoA) and LC (heptadecanoyl-CoA and stearoyl-CoA) substrates (**Fig. 1a,d**).

**Figure 1:**
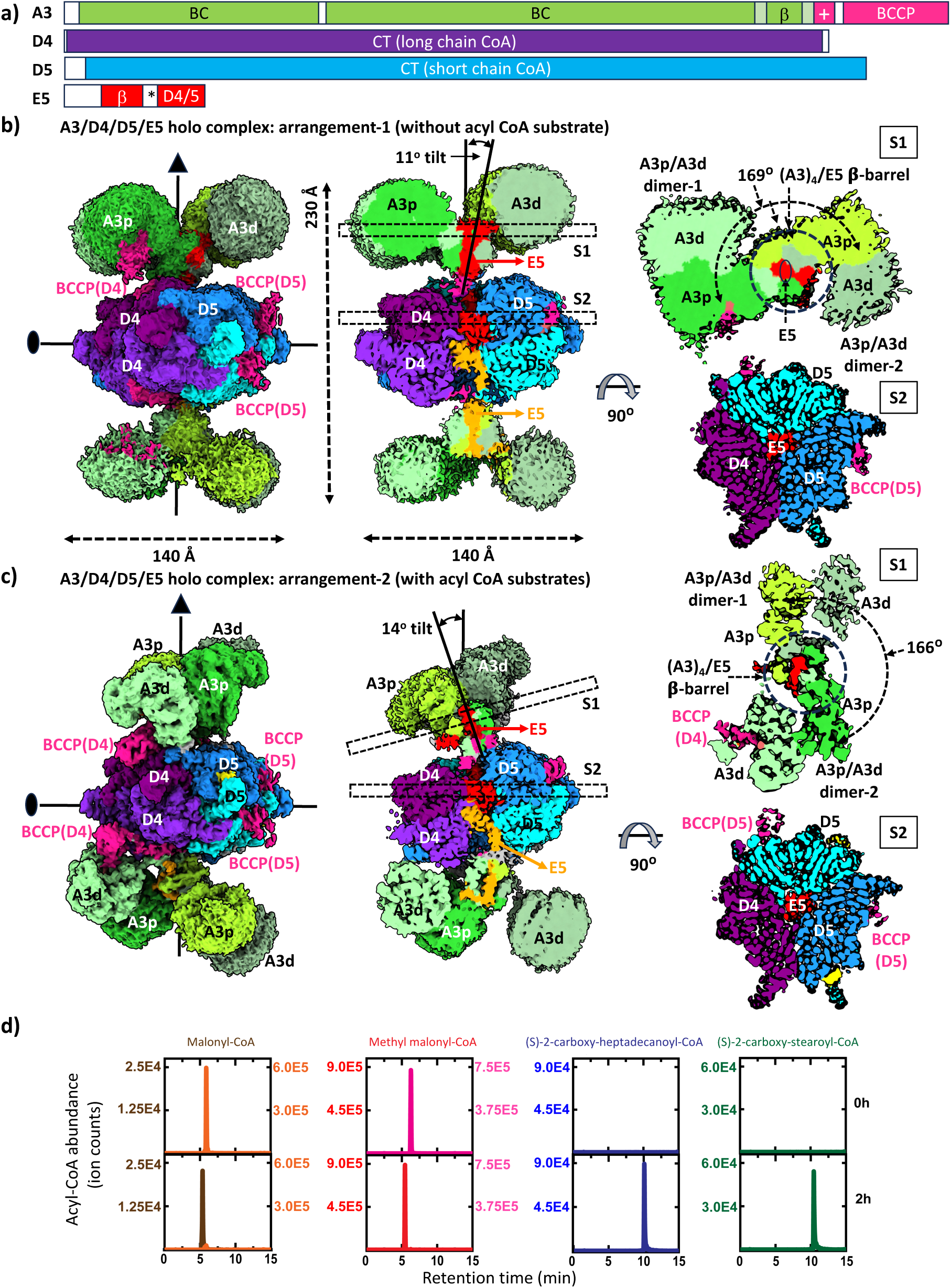
Overall arrangement of the mycobacterial AccA3/AccD4/AccD4/AccE holo complex with dual substrate specificity for LC and SC acyl-CoAs. a, domain structure of subunits participating in complex formation, proportional to sequence length. Colors of structurally visible segments: AccA3, green shades; AccA3 BCCP domains, magenta; AccD4, violet; AccD5, blue/cyan shades; AccE: orange/red. Abbreviations: β, β-barrel; +, positively charged BCCP linker; *, flexible linker in AccE5, connecting BC and CT assemblies; **b, c,** overall architecture of the AccA3/AccD4/AccD5/AccE5 holo complex without (b) and with (c) acyl-CoA substrates. Left, density presentation; central, sliced presentation across the center of the complex; right, sliced sections of (AccA3)_4_AccE5 BC (S1) and (AccD4)_2_/(AccD5)_4_/(AccE5)_2_ CT (S2) assemblies in top view. Structural elements are labeled and geometric parameters are indicated. Acronym shortcuts used: AccA3, A3; AccD4, D4; AccD5, D5, AccE5, E5. **d,** product formation from acetyl-CoA, propionyl-CoA, heptadecanoyl-CoA or stearoyl-CoA (left to right), demonstrating SC/LC acyl-CoA substrate activity. Corresponding product names are shown in colors matching the traces.

We then determined the high-resolution cryo-EM structures of this assembly, revealing a 16-subunit holoenzyme architecture with (AccA3)_4_–AccE5–[(AccD4)₂/(AccD5)₄]–AccE5– (AccA3)_4_ stoichiometry, both in the absence and presence of LC and SC acyl-CoA substrates (**Fig. 1b,c**). Its core is formed by a central hybrid CT assembly with a hexameric (AccD4)₂/(AccD5)₄ stoichiometry in a double-ring arrangement with 32-symmetry despite the mixed AccD4/AccD5 composition.

A unique feature of the complex is the critical anchoring and conformational flexibility of the AccE5 subunit dimer, which forms a central axis of this complex (**Fig. 1b**). Each AccE5 subunit is divided into an N-terminal α/β-hairpin motif and a small C-terminal α−helical hairpin domain, connected by a flexible linker (**Fig. 1a**, **Fig. 2c**). The AccE5 dimer places each of the two AccE5 subunits within one AccD4/(AccD5)_2_ layer of the central CT assembly to connect to the vis-à-vis AccA3 tetramer through its flexible linker. The AccE5 C-terminal hairpin domain mediates distinct interactions with each AccD4 subunit mediated through its C-terminal helix, while the remaining hairpin structure interacts with both AccD5 subunits (**Fig. 2a**). These data illustrate that the AccE5-mediated interactions are critical for forming the observed AccD4/(AccD5)_2_ hybrid assembly within each CT layer, as opposed to established homo-oligomeric CT assemblies without an AccE5-like subunit from reconstituted ACCase complexes (1, 7).

**Figure 2:**
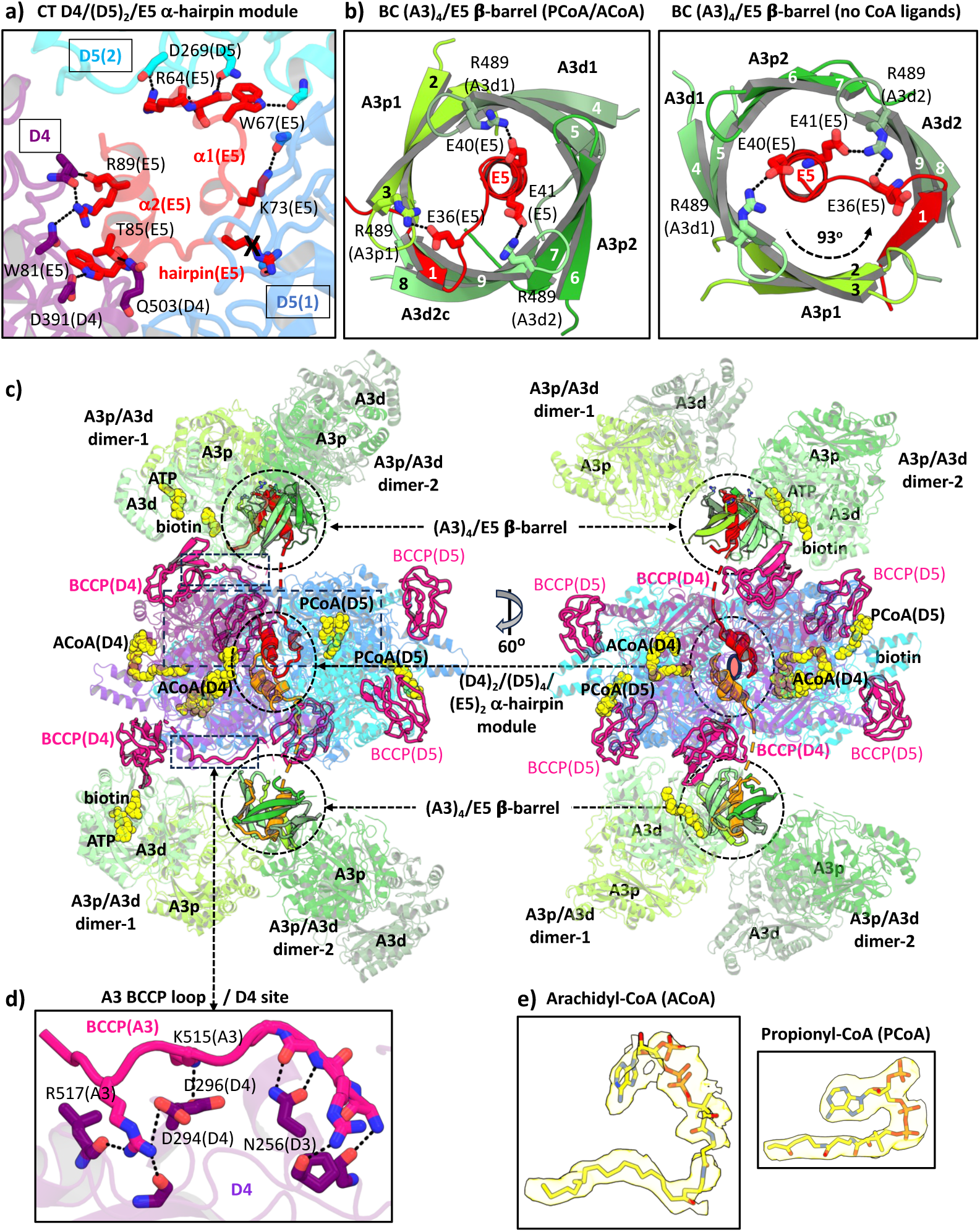
Structural elements that generate asymmetry, flexibility and multi-substrate specificity of the AccA3/AccD4/AccD4/AccE holo complex. **a**, CT AccD4/ AccD5/AccE5 α-hairpin module; **b,** BC (AccA3)_4_/AccE_5_ β-barrel in the presence (central) and absence (right) of acyl-CoA ligands, indicating a 93° rotation between the two states**. c,** cartoon presentation of the AccA3/AccD4/AccD4/AccE holo complex orientated as in Fig. 1 **(left)** and vertically rotated by 60°, allowing a view along the AccE5 dimeric twofold axis with the two AccD4 subunits of the hexameric CT complex in front. Bound ACoA and selected PCoA substrates (only when visible) are shown in yellow spheres; **d,** AccA3 BCCP loop / AccD4 interactions; **e**, ACoA and selected PCoA substrates within corresponding densities. All other conventions are as in Fig. 1.

The N-terminal AccE5 α/β-hairpin motif contributes to an unusual 9-stranded β−barrel by inserting a single β−strand into a fourfold repeated array of two-stranded β−hairpins from four AccA3 subunits, thus generating a central AccE5-mediated AccA3 tetramerization motif (**Fig. 2b middle and right panels)**. Due to the fourfold repeated β−sheet hairpin sequence of the AccA3 protomers, the same residue (R489) from three protomers forms equivalent salt bridges with three glutamate residues next to each other within the AccE5 α/β-hairpin sequence (E36, E40, E41), thus contributing to the structural integrity of the (AccA3)_4_/AccE5 β−barrel tetramerization motif. Although similar β-barrel motifs are found in single BC subunits of canonical single substrate ACCase complexes as well (2), in this structure it fundamentally differs by its composition from four AccA3 subunits and the observed asymmetry due to the additional AccE5 β−strand insertion.

Remarkably, the C-terminal helix of the AccE5 α/β-hairpin motif inserts into the central hole of this barrel, pointing its C-terminus followed by the flexible linker towards the vis-à-vis AccD4/(AccD5)_2_ layer of the central hybrid CT complex (**Fig. 1b middle panel, Fig. 2c**). Due to its uneven composition, the (AccA3)_4_/AccE5 β−barrel tilts by 11 degrees from the threefold axis of the central (AccD4)_2_/(AccD5)_4_ hybrid CT assembly. This asymmetry further propagates to the catalytic BC domains of the four AccA3 protomers that form two separate homodimers (**Fig. 1b, c middle and right panels, Fig. 2c**). In each dimer, one proximal protomer catalytic domain (AccA3p) directly interacts with central β−barrel residues of both AccA3 protomers (**Fig. 1b right panel, Fig. 2c**). The other protomer (AccA3d) is in a distal position, without any interactions with the central β−barrel. The two AccA3p/AccA3d dimers are related by uneven rotations around an axis established by the central (AccA3)_4_/AccE5 β−barrel. The larger rotation of 191 degrees is around the barrel segment with a single AccE5 β−strand inserted (**Fig. 1b right panel)**.

Finally, each of the AccA3 protomers comprises a C-terminal BCCP domain that transfers carboxybiotin from the AccA3 BC active site to one of the CT AccD4 and AccD5 active sites (**Fig. 1b left panel, Fig. 2c**). However, the resulting 8:6 stoichiometry of AccA3 and AccD4/AccD5 subunits in the holo complex generates a mismatch between eight AccA3-connected BCCP domains and six CT active sites. Of the six BCCP domains visible in this structure, four are bound to the four AccD5 CT active sites (**Fig. 2c**). Since the linkers to the remaining sequences of their parent AccA3 subunits are invisible, the precise connections are unknown. The other two visible BCCP domains, located at opposite faces of the central hexameric CT complex, are loosely connected to the surfaces of those donating AccA3p catalytic BC domains that are most tilted towards the two vis-à-vis AccD4 subunits of the central hexameric CT complex (**Fig. 1b, c left panel, Fig. 2c**). While there are no direct contacts between the BCCP domains and the AccD4 subunits, a highly positively charged segment of the connecting linker (residues 511-519, sequence RKKPKPRKR) directly interacts with several residues on each AccD4 surface (**Fig. 2c, d**).

Cryo-EM structures of the (AccA3)_8_/(AccD4)_2_/(AccA5)_4_/(AccE5)_2_ holo complexes in the presence of SC (propionyl-CoA, PCoA) or LC substrates (arachidic-CoA, ACoA) reveal that these substrates are specifically bound to the CT active site pockets of the AccD5 and AccD4 subunits, respectively (**Fig. 1c**, **Fig. 2c, e**). When comparing the three structures, we noticed a major conformational change in the holo complexes with bound acyl-CoA ligands, resulting in an even more tilted arrangement between the central hexameric CT assembly and the tetrameric BC assembly (**Fig. 1b, c, Fig. 2c**). This change is caused by a rotation of approximately 90 degrees of the central (AccA3)_4_/AccE5 tetramer β−barrel, which propagates into a corresponding rotation of the associated AccA3 catalytic domain dimers (**Fig. 1b, c right panels, Fig. 2b middle and right panels).** The rotation effectively switches the positions of neighboring AccA3 subunits and AccD4/AccD5 subunits facing each other (**Fig. 1b, c, Fig. 2c**). Therefore, the two visible BCCP domains vis-à-vis the two AccD4 subunits switch their origin from one of the AccA3p subunits to AccA3d subunits. In the presence of acyl-CoA ligands, the BCCP domain on its own also becomes associated with the surface of the AccD4 subunit (**Fig. 1c**, **Fig. 2c**). This observation suggests a role for all eight AccA3 subunits in BCCP-mediated carboxybiotin delivery, involving two AccA3 subunits for each of the two AccD4 CT subunit active sites.

In summary our data present a structure of a hybrid SC/LC ACCase complex that integrates two different CT subunits in a 4:2 stoichiometry. In addition, this complex is highly asymmetric and is suggestive of a rotational movement of the tetrameric BC assembly to be integral to its multifunctional catalytic mechanism. We expect these data to generate a plethora of functional and mechanistic research questions, allowing to substantially expanding our knowledge of hybrid and multifunctional enzyme complexes. Our data demonstrate the potential of extracting endogenous multi-protein complexes from their natural source as a superior alternative to the classical reconstitution of expressed protein components.

## Methods

### Bacterial strain and growth conditions

*M. smegmatis* mc^2^155 strain and the *M*. *smegmatis* Δ*accD1-*Δ*accA1* and Δ*accD2-*Δ*accA2* strains (1) were initially grown in a pre-culture using 7H9 medium supplemented with 10% ADS (BSA 50 g/L, glucose 20 g/L, and NaCl 8.1 g/L) for 64 hours at 37°C with shaking at 140 rpm. This pre-culture served as inoculum for a 2L culture. The main culture was grown in 7H9 broth supplemented with 0.2% glycerol, 0.1% glucose, and 0.05% Tween 80. Cultures were incubated at 37°C with continuous shaking at 140 rpm for 48 hours until reaching an optical density at 600 nm of 2.5. No antibiotics were added to the growth medium.

### Cell harvesting and lysis

Bacterial cells were harvested by centrifugation at 7,000 × g for 60 minutes at 20°C. The cell pellet was resuspended in lysis buffer containing 50 mM Tris-HCl (pH 7.5) and 300 mM NaCl. To prevent proteolysis, a protease inhibitor cocktail (Serva) was added to the resuspended cells. DNase I (1 mg/ml) was also included to reduce sample viscosity. Cells were lysed using a Sonoplus HD3200 sonicator at 45% amplitude for 6 minutes, using a 10-seconds on / 5-seconds off pulse cycle, while keeping the sample on ice. The cell lysate was clarified by centrifugation at 19,000 × g for 1 hour at 4°C, followed by filtration through a 0.45 µm filter.

### Protein purification

The filtered lysate from was incubated with pre-equilibrated streptavidin sepharose resin (Merck) for 2 hours at 4°C with gentle agitation. The resin was then extensively washed with lysis buffer to remove non-specifically bound proteins. The biotinylated protein complexes were eluted from the resin using 50 mM biotin in a buffer containing 50 mM Tris-HCl (pH 7.2) and 150 mM NaCl (**Extended Data Fig. 1a)**.

Elution fractions from strep-tactin affinity purification were analyzed by SDS-PAGE to assess protein purity. The pooled eluate was further purified by size exclusion chromatography using a Superose 6 10/300 column (GE Healthcare) equilibrated with 50 mM Tris-HCl (pH 7.5), 150 mM NaCl, and 2 mM DTT. Fractions containing the purified AccA3/AccD4/AccD5/AccE5 holo complex were pooled and concentrated using a centrifugal filter unit (**Extended Data Fig. 1b).**

### Mass spectrometry

Protein bands were excised from the gel and cut into 1 mm³ pieces for in-gel digestion as described (8). Gel pieces were dehydrated with 100% acetonitrile, reduced with 10 mM 1,4-dithiothreitol (DTT) for 30 min at 56°C, dehydrated again, and alkylated with 55 mM 2-chloroacetamide for 20 min at RT in the dark. Following a final dehydration, proteins were digested overnight at 37°C using 2 ng/µL trypsin in 50 mM ammonium bicarbonate. Peptides were extracted by sonication for 15 min, followed by centrifugation. A second extraction was performed by sonication for 15 min, using 50% acetonitrile and 1% formic acid at twice the gel volume. The pooled supernatants were dried by vacuum centrifugation and reconstituted in 4% acetonitrile with 1% formic acid for LC-MS/MS analysis. LC-MS/MS peptide analysis was performed on an UltiMate 3000 RSLCnano LC system (Thermo Fisher Scientific) equipped with a µ-Precolumn C18 PepMap™ 100, 300 µm i.d. × 5 mm, 5 µm, 100 Å trapping cartridge (Thermo Fisher Scientific) and an analytical column (nanoEase™ M/Z HSS T3, 75 µm i.d. × 250 mm, 1.8 µm, 100 Å) (Waters). Samples were trapped at 30 µL/min in 0.05% trifluoroacetic acid for 6 min. Peptides were eluted at 0.3 µL/min using a gradient of solvent B (3% DMSO, 0.1% formic acid in acetonitrile) against solvent A (3% DMSO, 0.1% formic acid in water). Peptides were introduced into an Orbitrap Fusion™ Lumos™ Tribrid™ mass spectrometer (Thermo Fisher Scientific) via a Pico-Tip emitter (360 µm OD × 20 µm ID, 10 µm tip; CoAnn Technologies) with a 2.2 kV spray voltage. The instrument operated in positive ion mode with a capillary temperature of 275 °C. Full MS scans were acquired in the Orbitrap profile mode from m/z 350–1,500 at 120,000 resolution (at m/z 200), with a 100 ms maximum injection time and standard AGC target. For MS/MS, the instrument operated in DDA mode, acquiring scans in the ion trap in rapid scan mode with a 35 ms maximum injection time and standard AGC target. Fragmentation was induced by HCD (30% normalized collision energy), and MS2 spectra were acquired in centroid mode. Database Search and AnalysisRaw files were converted to mzML format with MSConvert (ProteoWizard), applying peak picking for the 1000 most intense peaks, 64-bit encoding, and zlib compression. Files were searched using MSFragger in FragPipe (22.1-build02) against FASTA databases (MycolicibacteriumSmegmatis_ATCC700084_UP000000757_ID246196_entries6602_261020 22_dl11012023 and P4225 with common contaminants and reversed sequences). Search parameters included: carbamidomethylation (C, 57.0215) as a fixed modification; oxidation (M, 15.9949) and N-terminal acetylation (42.0106) as variable modifications. Mass error tolerances were 20 ppm (MS1) and 0.5 Da (MS2). Trypsin was specified as the protease, allowing a maximum of 2 missed cleavages and a minimum peptide length of 7. The FDR was set to 0.01 (1%) at both peptide and protein levels. The ‘Default’ FragPipe workflow was used with the following modifications: ionquant.maxlfq: 0, ionquant.mbr: 0, ionquant.minions: 2, ionquant.normalization: 0, ionquant.uniqueness: 1, msfragger.fragment_mass_tolerance: 0.5, msfragger.fragment_mass_units: 0, msfragger.misc.fragger.enzyme-dropdown-1: trypsin, msfragger.search_enzyme_name_1: trypsin, msfragger.search_enzyme_nocut_1: P, phi-report.filter: --sequential --prot 0.01 -- razor, quantitation.run-label-free-quant: true.

### Enzymatic activity assays

The assay reaction mix contained 50 mM HEPES (pH 7.5), 5 mM Mg-ATP, 5 mM NaHCO_3_, and 100 µM of acyl-CoA substrates (acetyl-CoA, propionyl-CoA, heptadecanoyl-CoA or stearoyl-CoA, separate reaction for each substrate). The reaction was initiated by adding 10 µM of purified AccA3/AccD4/AccD5/AccE5 holo complex. After 2 hours of incubation, the reaction aliquots were quenched with cold acidic acetonitrile (0.2% acetic acid). The control reactions were done either without ATP or without AccA3/AccD4/AccD5/AccE5 holo complex. All experiments were carried out at 25 °C.

The formation of the carboxylated products was detected using liquid chromatography-mass spectrometry (LC-MS). For LC, an Infinity 1290 II LC-MS system (Agilent) was utilized.

Substrates and products were separated using a 150 mm × 2.1 mm Cogent 4 Diamond Hydride^TM^ HPLC column (MicroSolv Technology). The gradient used was defined according to an established protocol with additional equilibration at the end (9). Solvent A was 0.2% acetic acid in water and solvent B was 0.2% acetic acid in acetonitrile. The gradient (% solvent B) used was: 0–2 min, 85%; 3–5 min, 80%; 6–7 min, 75%; 8–9 min, 70%; 10–11.1 min, 50%; 11.1–14 min, 20%; 14.1–18 min 5%, followed by a 1 min re-equilibration period at 85% B at a flow rate of 0.4 ml/min. The column temperature was kept constant at 20 °C.

The HPLC system was coupled with an Accurate Mass 6230 TOF apparatus (Agilent) for metabolite identification. To achieve dynamic mass axis calibration, a reference mass solution was continuously infused using an isocratic pump with a 100:1 splitter. ESI capillary was set at 3500 V and fragmentor was set at 110 V. The nebulizer pressure was set to 35 psi, while the nitrogen drying gas was delivered at a flow rate of 13 L/min and maintained at a temperature of 250 °C. The sheath gas was set to a temperature of 350 °C with a flow rate of 12 L/min to support efficient desolvation. Mass spectrometric data were acquired at a rate of 1 spectrum per second across an m/z range of 50–1200. The instrument consistently provided accurate mass measurements with a mass error below 5 ppm, a resolution ranging from 10,000 to 25,000 over the m/z range of 121–955, and a dynamic range spanning five orders of magnitude. Data were acquired in centroid mode using the 4 GHz (extended dynamic range) setting. The data were analyzed with the MassHunter Qualitative Analysis software (Agilent). A mass tolerance of <0.005 Da was applied for metabolite identities.

### Cryo-EM data acquisition, processing and 3D reconstruction

For structure determination of the AccA3/AccD4/AccD5/AccE5 holo complex, 3 µL of sample at a concentration of 4 mg/ml was applied to glow-discharged Quantifoil R2/1 300-mesh copper grids. The grids were blotted for 2 seconds at 4°C and 100% humidity before being plunge-frozen in a liquid ethane-propane mix using a Vitrobot Mark IV (Thermo Fisher Scientific). For samples in the presence of acyl-CoA substrates, cryo-EM grids were prepared by mixing AccA3/AccD4/AccD5/AccE5 holo complex at concentration of 5 mg/ml with substrates to a final concentrations of 1 mM propionyl-CoA (Merck) or 1 mM arachidyl-CoA (Avanti Lipids), 4 mM ATP, 20 mM NaHCO₃, and 5 mM MgCl₂. Acyl-CoA substrates were added from 10 mM stocks and other substrates (ATP, NaHCO₃, and MgCl₂) were added from 100 mM stocks. R2/1 Quantifoil grids were glow-discharged for 120 s. The substrates were mixed immediately before being applied to the grids, the total time from mixing to plunge-freezing in the liquid ethane-propane mix was 45 seconds. The grids were blotted for 2.5 s with a blot force of −7 at 4°C and 100% humidity using a Vitrobot Mark IV (Thermo Fisher Scientific).

Cryo-EM data for all samples were collected on a Titan Krios G3 microscope (Thermo Fisher Scientific) operating at 300 kV, equipped with a K3 direct electron detector (Gatan) and a Quantum energy filter (20 eV slit width) operating in counting mode. Data were recorded at a nominal magnification of 130,000x, corresponding to a calibrated pixel size of 0.68 Å/pixel. In total, 18,636 movies were recorded for AccA3/AccD4/AccD5/AccE5 holo complex in the absence of acyl-CoA ligands, 10,897 in the presence of propionyl-CoA, and 6,272 in the presence of arachidyl-CoA. Each movie consisted of 40 frames, with a total exposure of 44 e⁻/Å² and a defocus range of −0.5 to −2.25 µm. The EPU program (Thermo Fisher Scientific) was used for automated data acquisition.

All datasets were processed in cryoSPARC v4.2 (10). Movie frames were motion-corrected and dose-weighted using MotionCor2, and the contrast transfer function (CTF) was estimated using CTFFIND4, both implemented within cryoSPARC. Particles were picked using a combination of blob and template pickers, extracted with a 560-pixel box size and binned 2x, and subjected to multiple rounds of 2D classification.

Particles from the best 2D classes were selected to generate an initial 3D model. After several rounds of heterogeneous refinement, a subset of particles was selected for a final refinement and re-extracted with a 560-pixel box size. The refinement yielded 3D reconstructions at a global resolution of 2.4 Å in the absence of any acyl-CoA ligands, the presence of propionyl-CoA, and the presence of arachidyl-CoA, as determined by the gold-standard Fourier shell correlation (FSC) 0.143 criterion. The resolution achieved mainly arises from the central (AccD4)_2_/(AccD5)_4_/(AccE5)_2_ core **(Extended Data Fig. 2, 3, 4).**

In the global map of the structure of the AccA3/AccD4/AccD5/AccE5 holo complex without acyl-CoA substrate, the electron density for both peripheral (AccA3)_4_AccE5 assemblies was significantly weaker than for the central (AccD4)_2_/(AccD5)_4_/(AccE5)_2_ core **(Extended Data Fig. 2)**. In the structure of the AccA3/AccD4/AccD5/AccE5 holo complex in the presence of propionyl-CoA, one 3D class showed well-defined density for the complete complex **(Extended Data Fig. 3).** For the structure of the AccA3/AccD4/AccD5/AccE5 holo complex in the presence of arachidyl-CoA structure, one (AccA3)_4_AccE5 assembly was well-resolved, while the other one was only poorly resolved **(Extended Data Fig. 4)**. To improve the density in this region, focused local refinement was performed for all structures using a focused mask on the AccA3 region. This resulted in locally refined maps of one of the two (AccA3)_4_AccE5 assemblies at 3.2 Å (no acyl-CoA ligands), 3.0 Å (propionyl-CoA), and 3.5 Å (arachidyl-CoA). For the second (AccA3)_4_AccE5 assembly, a resolution of 3.3 Å was achieved for the structure in the presence propionyl-CoA (**Extended Data Fig. 2, 3, 4**), whereas local refinement did not yield well-defined densities for the corresponding assemblies in the absence of acyl-CoA ligands and in the presences of arachidyl-CoA structure.

### Model building, refinement, validation and analysis

An initial model of a AccD5 hexamer was predicted by AlphaFold 3 (11) was fitted into the cryo-EM density map of the AccA3/AccD4/AccD5/AccE5 holo complex in the absence of acyl-CoA ligands, using ChimeraX (12). The sequences for AccD4 and the C-terminus of AccE5 were subsequently assigned based on high-resolution features in the map, allowing model building of the complete (AccD4)_2_/(AccD5)_4_/(AccE5)_2_ core assembly. During this procedure two AccD5 subunits were replaced by AccD4 subunits, thus generating a hybrid CT core complex with 4:2 AccD5/AccD4 stoichiometry. Iterative refinement was performed using Servalcat (13) within the CCPEM Doppio suite. Rotamers and outliers were corrected in Isolde (14) within ChimeraX and manually adjusted in Coot (15).

For the AccA3/AccE5 assembly, an AlphaFold 3 model was used as the initial template. However, since the EM map considerably deviated from the predicted symmetric AccA3 tetrameric arrangement, extensive manual remodeling and refinement was required, using Coot (15) and Servalcat (13), respectively.

For the propionyl-CoA and arachidyl-CoA bound structures, the refined structure of the AccA3/AccD4/AccD5/AccE5 holo complex in the absence of acyl-CoA ligands was used as a template. While the central (AccD4)_2_/(AccD5)_4_/(AccE5)_2_ core assembly fitted well into the corresponding maps, the (AccA3)4/AccE5 assembly did not, indicating a different conformation triggered by the presence of these acyl-CoA substrates. These regions were manually rebuilt in Coot, corrected for rotamer and Ramachandran outliers in Coot and Isolde, and further refined in Servalcat.

While the AccA3/AccD4/AccD5/AccE5 holo complex structures in the absence of acyl-CoA ligands and in the presence of arachidyl-CoA showed high-resolution density for only one of the two (AccA3)₄/AccE₅ tetramers, the propionyl-CoA structure revealed both flanking (AccA3)₄/AccE₅ assemblies at sufficient resolution for molecular interpretation. Therefore, the complete (AccA3)₈/(AccD4)₂/(AccD5)₄/(AccE5)₂ holo complex in the presence of propionyl-CoA structure and truncated (AccA3)₄/(AccD4)₂/(AccD5)₄/(AccE5)₂ complexes in the absence of acyl-CoA ligands and in the presence of arachidyl-CoA were refined and used for further analysis.

Sequence confidence was validated using the CheckMySequence program (16) in the CCPEM Doppio suite. The final models were validated using the MolProbity server, also implemented in CCPEM Doppio. Geometry analysis was carried out in Pymol (17), using PSICO(18) and Draw_Rotation_Axis plugins(19).

## Acknowledgements

We thank the CSSB cryo-EM facility for microscope access and technical support. We acknowledge the EMBL Heidelberg Proteomics Core Facility for performing the mass spectrometry analysis.

## Author Contributions

E.M. and M.W. designed the project. E.M. and H.M.T. performed the experimental work. All authors analyzed the data. E.M. and M.W. wrote the paper. L.P.S.d.C. and M.W. supported the project.

## References

1. Ehebauer MT, Zimmermann M, Jakobi AJ, Noens EE, Laubitz D, Cichocki B, et al. Characterization of the mycobacterial acyl-CoA carboxylase holo complexes reveals their functional expansion into amino acid catabolism. PLoS Pathog. 2015;11(2):e1004623.

2. Tong L. Structure and function of biotin-dependent carboxylases. Cell Mol Life Sci. 2013;70(5):863–91.

3. Reddy MC, Breda A, Bruning JB, Sherekar M, Valluru S, Thurman C, et al. Structure, activity, and inhibition of the Carboxyltransferase beta-subunit of acetyl coenzyme A carboxylase (AccD6) from Mycobacterium tuberculosis. Antimicrob Agents Chemother. 2014;58(10):6122–32.

4. North EJ, Jackson M, Lee RE. New approaches to target the mycolic acid biosynthesis pathway for the development of tuberculosis therapeutics. Curr Pharm Des. 2014;20(27):4357–78.

5. Bazet Lyonnet B, Diacovich L, Gago G, Spina L, Bardou F, Lemassu A, et al. Functional reconstitution of the Mycobacterium tuberculosis long-chain acyl-CoA carboxylase from multiple acyl-CoA subunits. FEBS J. 2017;284(7):1110–25.

6. Oh TJ, Daniel J, Kim HJ, Sirakova TD, Kolattukudy PE. Identification and characterization of Rv3281 as a novel subunit of a biotin-dependent acyl-CoA Carboxylase in Mycobacterium tuberculosis H37Rv. J Biol Chem. 2006;281(7):3899–908.

7. Gago G, Kurth D, Diacovich L, Tsai SC, Gramajo H. Biochemical and structural characterization of an essential acyl coenzyme A carboxylase from Mycobacterium tuberculosis. J Bacteriol. 2006;188(2):477–86.

8. Shevchenko A, Tomas H, Havli J, Olsen JV, Mann M. In-gel digestion for mass spectrometric characterization of proteins and proteomes. Nature protocols. 2006;1(6):2856–60.

9. Pesek JJ, Matyska MT, Watanabe S, Makhanov M, Lopez A, Alejo K, et al. Evaluation of silica hydride materials for the LC–MS analysis of cathinones and benzylpiperazines. Forensic Chemistry. 2018;8:90–4.

10. Punjani A, Rubinstein JL, Fleet DJ, Brubaker MA. cryoSPARC: algorithms for rapid unsupervised cryo-EM structure determination. Nat Methods. 2017;14(3):290–6.

11. Abramson J, Adler J, Dunger J, Evans R, Green T, Pritzel A, et al. Accurate structure prediction of biomolecular interactions with AlphaFold 3. Nature. 2024;630(8016):493–500.

12. Pettersen EF, Goddard TD, Huang CC, Meng EC, Couch GS, Croll TI, et al. UCSF ChimeraX: Structure visualization for researchers, educators, and developers. Protein Sci. 2021;30(1):70–82.

13. Yamashita K, Palmer CM, Burnley T, Murshudov GN. Cryo-EM single-particle structure refinement and map calculation using Servalcat. Acta Crystallogr D Struct Biol. 2021;77(Pt 10):1282–91.

14. Croll TI. ISOLDE: a physically realistic environment for model building into low-resolution electron-density maps. Acta Crystallogr D Struct Biol. 2018;74(Pt 6):519–30.

15. Casanal A, Lohkamp B, Emsley P. Current developments in Coot for macromolecular model building of Electron Cryo-microscopy and Crystallographic Data. Protein Sci. 2020;29(4):1069–78.

16. Chojnowski G. Sequence-assignment validation in cryo-EM models with checkMySequence. Acta Crystallogr D Struct Biol. 2022;78(Pt 7):806–16.

17. Schrödinger L. The PyMOL Molecular Graphics System, Version 3.1. Schrödinger, LLC. 2025.

18. Holder T. pymol-psico: PyMOL Script Collection. GitHub. 2023.

19. Calvo PG. draw_rotation_axis.py. GitHub. 2014.

