## Supplementary Data for "Architecture of an asymmetric mycobacterial short chain/long chain acyl-CoA carboxylase"

### Extended Data

#### Extended Data Table 1. Cryo-EM data collection, refinement and validation statistics

| | A3/D4/D5/E5 #1 | $(A3)_4$/E5 #2 | A3/D4/D5/E5 + PCoA #3 | A3/D4/D5/E5 + ACoA #4 |
|---|---|---|---|---|
| **Data collection and processing** | | | | |
| Magnification | 130,000 | | 130,000 | 130,000 |
| Voltage (kV) | 300 | | 300 | 300 |
| Detector / energy filter | K3/20 eV | | K3/20 eV | K3/20 eV |
| Electron exposure (e–/Å$^2$) | 44.4 | | 44.1 | 44.0 |
| Initial particle images (no.) | 1,279,569 | | 639,8521 | 353,856 |
| Final particle images (no.) | 251,974 | | 52,469 | 76,111 |
| Map resolution (Å) | 2.4 | 3.2 | 2.4 | 2.4 |
| **Refinement** | | | | |
| Initial model used | AF3 model | | #1 | #3 |
| Model resolution (Å) | 2.3 | 3.3 | 2.4 | 2.4 |
| CC (mask) | 0.90 | 0.86 | 0.85 | 0.81 |
| Model composition | | | | |
| Non-hydrogen atoms | 42,209 | 15,386 | 57,621 | 40,162 |
| Protein residues | 5,563 | 2,034 | 7,565 | 5,253 |
| Ligands | 5 | 1 | 11 | 4 |
| *B* factors (Å$^2$) | | | | |
| Protein | 81 | 80 | 56 | 58 |
| Ligand | 132 | 52 | 102 | 179 |
| R.m.s. deviations | | | | |
| Bond lengths (Å) | 0.0116 | 0.0106 | 0.0123 | 0.01 |
| Bond angles (°) | 1.82 | 1.67 | 1.79 | 1.65 |
| Validation | | | | |
| MolProbity score | 0.91 | 1.13 | 0.79 | 0.86 |
| Clashscore | 0.58 | 1.34 | 0.37 | 1.0 |
| Poor rotamers (%) | 0.23 | 0.32 | 0.63 | 0.46 |
| Ramachandran plot | | | | |
| Favored (%) | 96.65 | 96.04 | 97.14 | 97.70 |
| Allowed (%) | 3.19 | 3.86 | 2.63 | 2.22 |
| Disallowed (%) | 0.16 | 0.10 | 0.23 | 0.08 |

**Extended Data Table 2. LC–MS/MS identification of ACCase subunits**

| Protein identified | Molecular weight (kDa) | Total intensity | Max HyperScore | Coverage (%) | Total intensity | Max HyperScore | Coverage (%) |
|---|---|---|---|---|---|---|---|
| | | *M. smegmatis* mc$^2$155 | | | *M. smegmatis* Δ*accD1*-Δ*accA1*, Δ*accD2*-Δ*accA2* | | |
| AccA3 | 63.1 | 4.59E+10 | 85.5 | 91.0 | 3.22E+10 | 83.9 | 78.3 |
| AccD5 | 58.4 | 9.35E+10 | 79.7 | 71.8 | 7.29E+10 | 71.3 | 71.0 |
| AccD4 | 56.2 | 5.73E+10 | 82.2 | 83.4 | 5.07E+09 | 73.0 | 82.0 |
| AccE5 | 10.3 | 2.03E+09 | 30.5 | 63.8 | 1.09E+09 | 37.8 | 63.8 |
| AccA1 | 70.0 | 6.28E+05 | — | — | — | — | — |
| AccD1 | 54.8 | 2.37E+08 | 66.5 | 57.7 | — | — | — |
| AccD2 | 56.6 | 3.95E+07 | 56.4 | 37.1 | — | — | — |

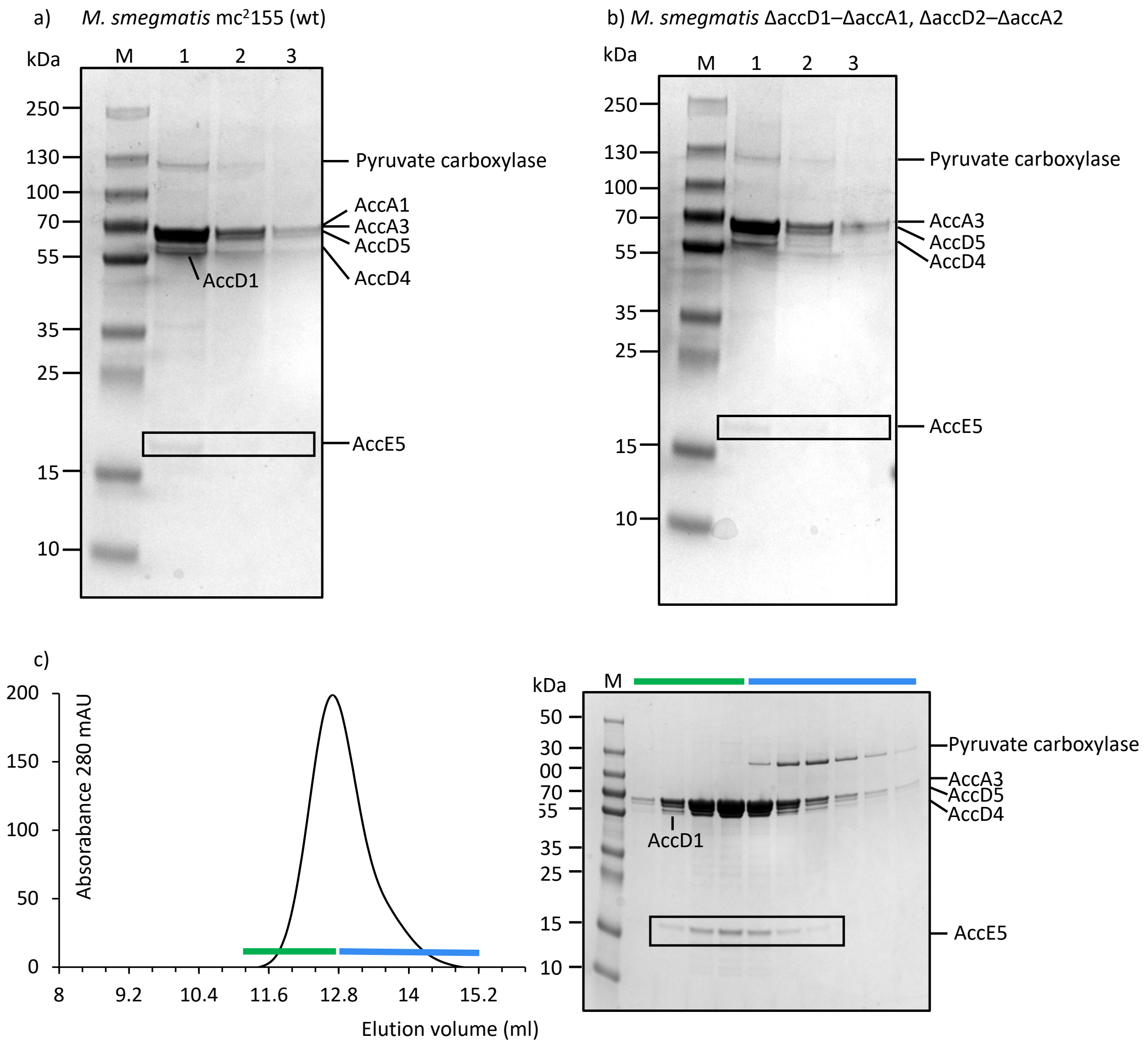


**Extended Data Figure 1: Endogenous purification of biotinylated proteins using streptavidin resin from *M. smegmatis*. a,** Strep-Tactin affinity purification of biotinylated proteins from cell lysates of *M. smegmatis* mc$^2$155 and *M. smegmatis ΔaccD1–ΔaccA1* and *ΔaccD2–ΔaccA2* strains. SDS–PAGE of fractions eluted with 50 mM biotin. Lanes M: molecular-weight marker; lanes 1–3: sequential elutions with 50 mM biotin. Bands identified by mass spectrometry correspond to pyruvate carboxylase (130 kDa), AccA1/AccA3 (70 kDa), AccD1/AccD4/AccD5 (60 kDa), and AccE5 (17 kDa). **b**, size-exclusion chromatography profile of the purified AccA3/AccD4/AccD5/AccE5 holo complex and SDS–PAGE of fractions across the main elution peak. Fractions marked by the green bar contain AccA1, AccA3, AccD4,

AccD5, AccD1 and AccE5; the blue bar indicates fractions also containing pyruvate carboxylase.

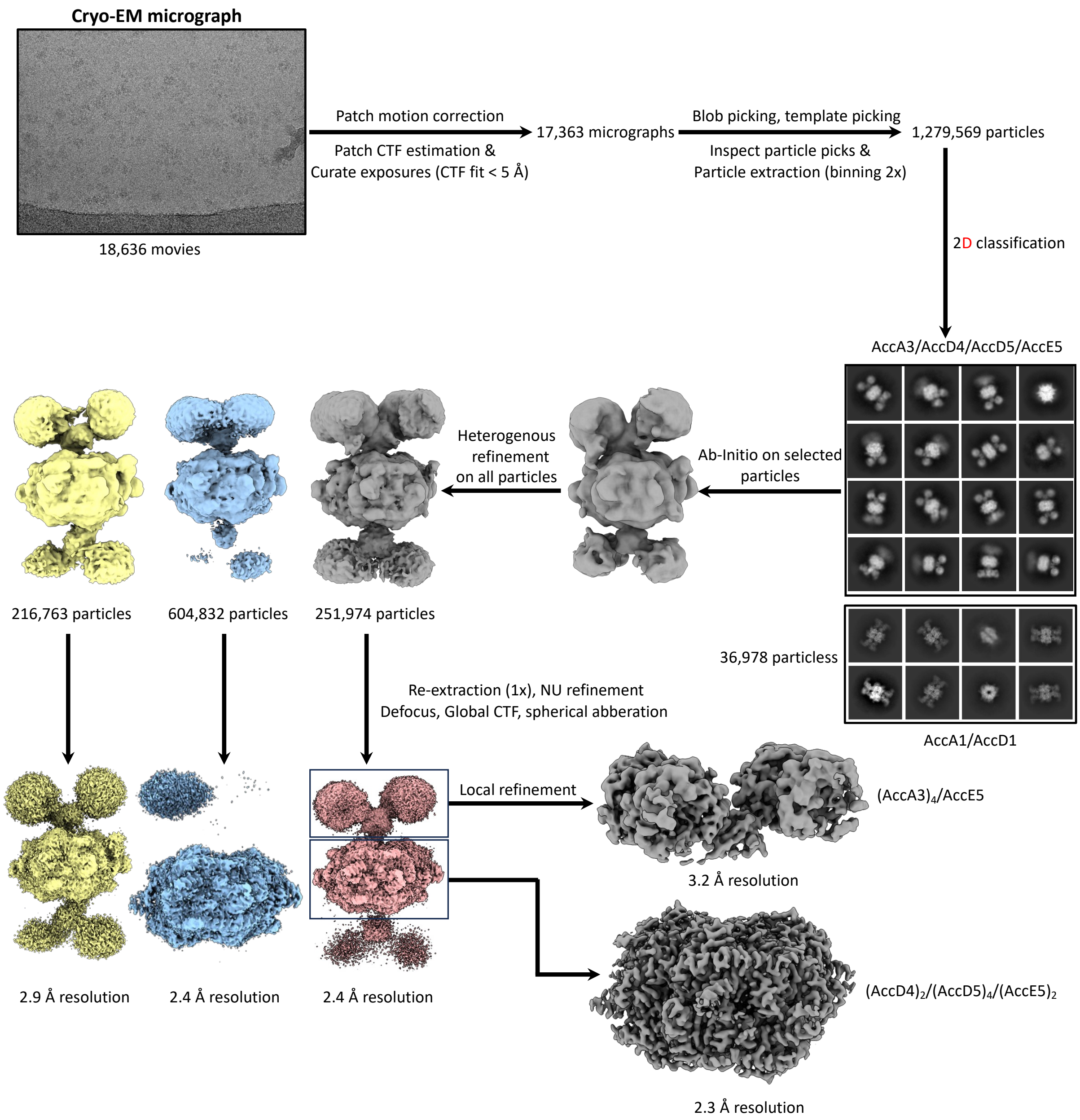


**Extended Data Figure 2: Cryo-EM data processing workflow for AccA3/AccD4/AccD5/AccE5 holo complex**. Schematic of the cryo-EM data processing pipeline, performed in cryoSPARC. A total of 18,636 movies was processed, yielding 1,279,569 particles. After 2D classification and several rounds of *ab initio* and heterogeneous refinement, a subset of 251,974 particles was selected. This particle set was subjected to re-extraction and non-uniform (NU) refinement, resulting in a 2.4 Å resolution map. Subsequent local refinement resolved the (AccD4)$_2$/(AccD5)$_4$/(AccE5)$_2$ core to 2.3 Å resolution and one of the two (AccA3)$_2$/AccE5 segments to 3.2 Å resolution. Composite maps were subsequently generated in ChimeraX for model building and refinement.

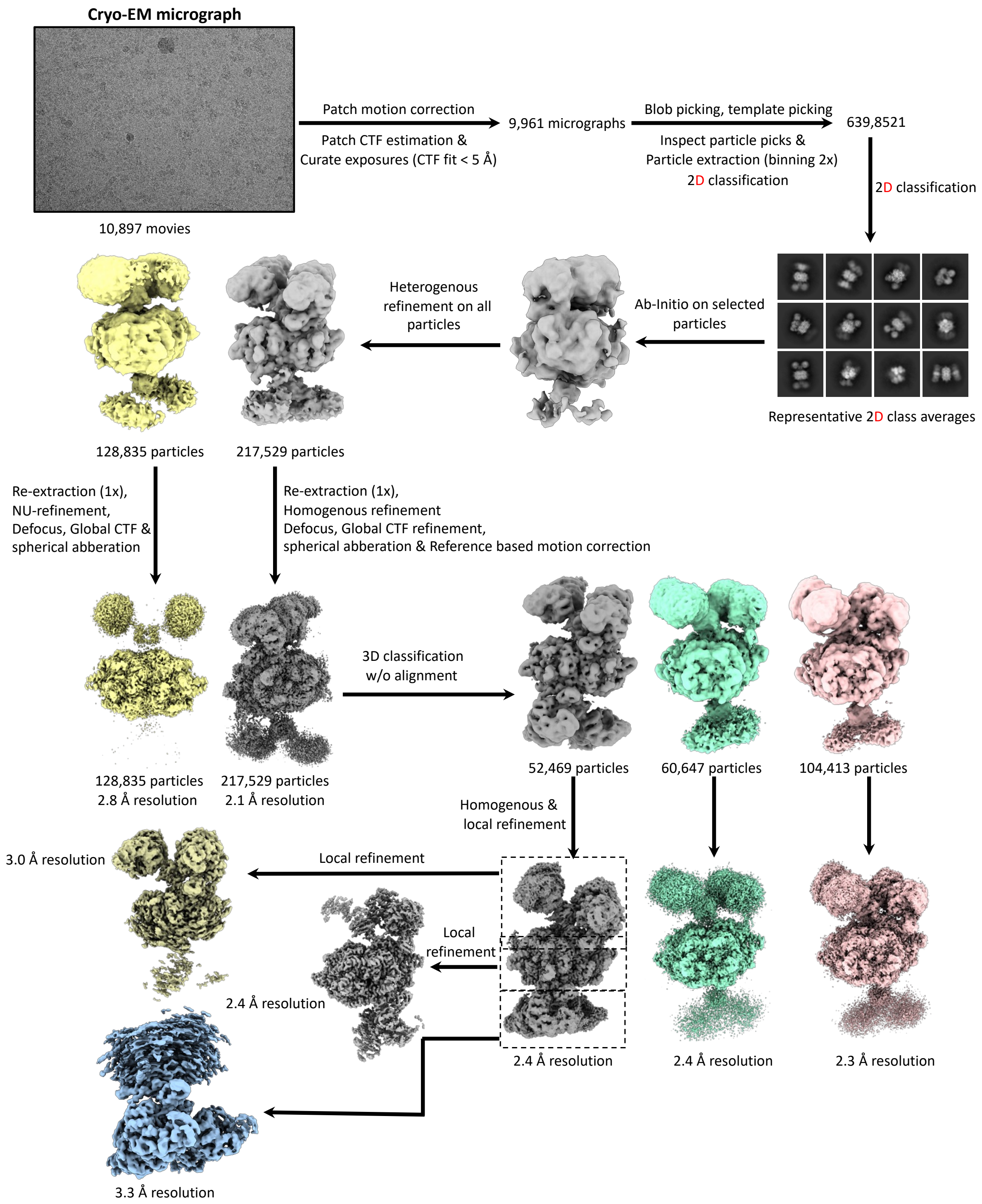


**Extended Data Figure 3: Cryo-EM data processing workflow for AccA3/AccD4/AccD5/AccE5 complex with propionyl-CoA.** A dataset of 10,897 movies was processed. Several rounds of 2D-classification, *ab initio* and heterogeneous refinement were performed to obtain a homogenous set of particles. Two classes corresponding to particle sets of 128,835 and 217,529 were selected for homogenous refinement, which yielded resolutions of 2.8 and 2.1 Å, respectively. Maps corresponding to 217,529 particles were sorted using 3D classification without alignment. This sorting resulted in multiple subsets of 52,469, 60,647 and 104,413 particles, which were then subjected to homogenous refinement that yielded maps with resolutions between 2.3 and 2.4 Å . The class that displayed the complete complex was used for local refinements focusing on the upper $(AccA3)_2$/AccE5 region, central $(AccD4)_2/(AccD5)_4/(AccE5)_2$ core and the bottom $(AccA3)_2$/AccE5 region that yielded resolutions of 3.0, 2.4 and 3.3 Å, repectively.

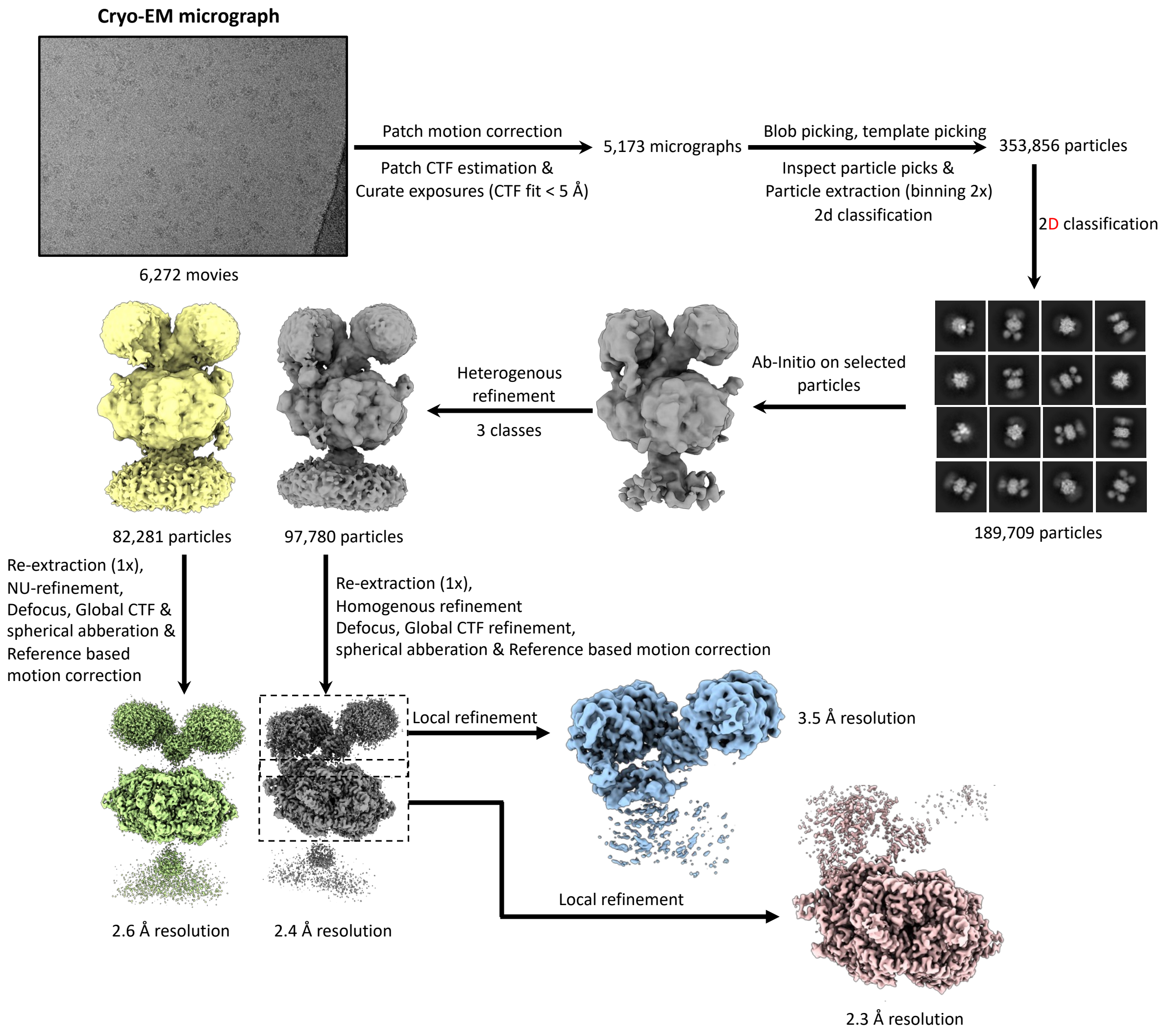


**Extended Data Figure 4: Cryo-EM data processing workflow for AccA3/AccD4/AccD5/AccE5 complex with Arachidyl-CoA.** From 6,272 movies, an initial set of 353,856 particles was picked. After 2D classification, 189,709 particles were used for heterogeneous refinement to generate three classes. The resulting particle subsets 82,281 and 97,780 particles were further refined yielding 2.6 Å and 2.4 Å maps. The class with 82, 281 particels was used for local refinement one of the two $(AccA3)_2$/AccE5 region and central $(AccD4)_2/(AccD5)_4/(AccE5)_2$ core of the complex, yielding final maps at 3.5 Å and 2.4 Å resolution.

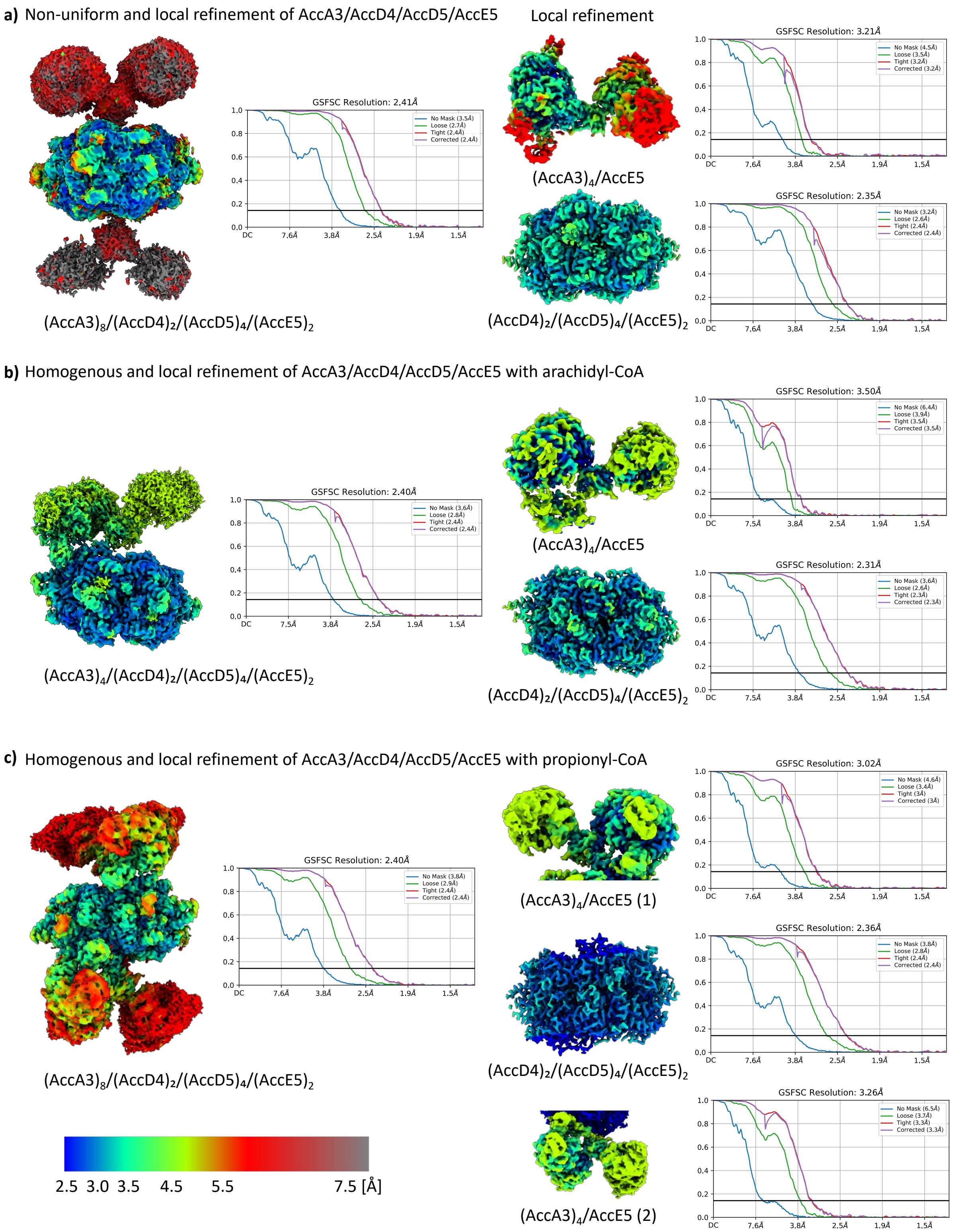


**Extended Data Figure 5: Local resolution estimation of AccA3/AccD4/AccD5/AccE5 complex without and with acyl-CoA substrates. a,** non-uniform refinement of AccA3/AccD4/AccD5/AccE5 holo complex without acyl-CoA substrates. The overall map is at resolution 2.4 Å. Subsequent local refinement of this map resolved one of the two $(AccA3)_2$/AccE5 region to 3.2 Å and the $(AccD4)_2/(AccD5)_4/(AccE5)_2$ core to 2.4 Å resolution. **b,** homogenous refinement of AccA3/AccD4/AccD5/AccE5 in the presence of arachidyl-CoA. The overall map is at 2.4 Å resolution. Local refinement resolved the upper region to 3.5 Å and the core to 2.3 Å resolution. **c,** homogenous refinement of AccA3/AccD4/AccD5/AccE5 with propionyl-CoA: The overall map is at 2.4 Å resolution. Further 3D classification and local refinement yielded maps with resolutions of 3.0 Å , 2.4 Å , and 3.3 Å. The maps are colored according to local resolution (in Å), as indicated by a color bar. The GSFSC plots show the correlation between two independent half-maps, with the final resolution determined by the 0.143 FSC criterion (horizontal black line).
